# Cadherin-11 integrates Piezo1 and interleukin-6 signaling to promote fibroblast activation

**DOI:** 10.1101/2025.09.16.674909

**Authors:** Leilani R. Astrab, Steven R. Caliari

## Abstract

Persistent fibroblast activation drives tissue fibrosis, yet how mechanical and inflammatory cues are integrated to promote this aberrant behavior remains unclear. Using a hyaluronic acid (HA)-based hydrogel platform to model normal and fibrotic lung mechanics, we examine the roles of Piezo1 and cadherin-11 (CDH11), both implicated in M2 macrophage-fibroblast crosstalk during pulmonary fibrosis progression, in interleukin (IL)-6-mediated fibroblast activation. While both Piezo1 and CDH11 expression increase in activated fibroblasts, blocking IL-6 signaling decreases CDH11, but not Piezo1, expression. Instead, Piezo1 activity promotes nuclear accumulation of the calcium-dependent transcription factor NFAT1. While Piezo1 inhibition moderately reduces CDH11 expression, it does not prevent fibroblast activation as measured by spreading and type I collagen expression, whereas CDH11 knockout suppresses fibroblast activation metrics, reduces Piezo1 expression, and decreases IL-6 secretion in both fibroblast only and fibroblast-M2 macrophage co-cultures. Furthermore, CDH11 levels increase in parallel with progressive fibroblast activation, highlighting its role in promoting this pro-fibrotic phenotype. Together, these findings underscore a previously unrecognized signaling axis in which CDH11 serves as a key mediator of sustained fibroblast activation, coordinating mechanical and inflammatory cues, and highlight CDH11 as a potential therapeutic target in pulmonary fibrosis.

## INTRODUCTION

Pulmonary fibrosis is a pathological wound healing process characterized by thickening of the alveolar interstitial tissue, which impairs lung function and can ultimately lead to organ failure.^1^ The most common form, idiopathic pulmonary fibrosis (IPF), has no known cause and currently lacks effective treatments to stop or reverse disease progression.^2–4^ However, it has been established that fibroblasts, macrophages, and the extracellular matrix (ECM) play central roles in fibrogenesis.^5–11^ These key contributors have dynamic and reciprocal relationships, where macrophages secrete cytokines that activate fibroblasts, fibroblasts activate to ECM-producing myofibroblasts, and both fibroblasts and macrophages respond to mechanical cues from the ECM through mechanotransduction.^5–11^ These interconnected pathways warrant the need for a more comprehensive understanding of the cell-cell and cell-matrix interactions contributing to IPF progression in order to develop more effective therapeutics.

While macrophages are highly plastic cells capable of adopting a broad spectrum of phenotypes essential for orchestrating proper wound healing and tissue repair, the alternatively activated “M2” macrophage subset has been implicated in fibrosis progression.^5–7,12,13^ In normal wound healing, M2 macrophages promote tissue remodeling and resolution by activating fibroblasts to produce ECM that helps restore tissue integrity.^14,15^ However, when wound healing is dysregulated in diseases like IPF, these same interactions lead to pathological scarring.^14,15^ Recent work from our lab showed that interleukin-6 (IL-6) is one of the mediators driving M2 macrophage-induced fibroblast activation.^16^ In addition to other studies that have demonstrated the pro-fibrotic effect of IL-6 both *in vitro* and *in vivo*, these findings highlight a crucial signaling pathway through which immune-stromal interactions contribute to fibrosis progression.^17–21^ Furthermore, our work demonstrated that M2 macrophage crosstalk is capable of activating fibroblasts even on soft viscoelastic hydrogels mimicking normal tissue mechanics, and that blocking IL-6 signaling disrupts this activation process.^16^

In addition to soluble signals like IL-6, direct mediators of M2 macrophage-fibroblast crosstalk have also been connected to fibroblast activation in IPF. These mediators include the transmembrane protein cadherin-11 (CDH11) and the stretch-activated membrane channel (SAC) Piezo1.^22–27^ CDH11 is upregulated in IPF, where it sustains M2 macrophage–fibroblast communication and promotes a profibrotic environment by driving persistent transforming growth factor beta (TGF-β) signaling.^22–24^ Recent findings have also identified Piezo1 as an early mediator of M2 macrophage–fibroblast crosstalk, acting through αvβ3 integrin engagement to initiate fibroblast activation.^25^ While both CDH11 and Piezo1 have been implicated in macrophage-fibroblast crosstalk, they also respond to cues from the surrounding ECM environment. Notably, CDH11 is the only cadherin shown to interact with the ECM, through its binding partner syndecan-4, which binds to fibronectin and links cell adhesion directly to matrix interactions.^28,29^ The role of Piezo1 in mechanosensing has been well-established, with studies demonstrating its role in focal adhesion formation and cytoskeletal remodeling to regulate cellular responses to substrate mechanics.^30–32^ Although both CDH11 and Piezo1 have been independently linked to promoting IL-6 signaling in fibrosis progression, their roles in pulmonary fibrosis have not been investigated.^33–37^ In this context, the ability of CDH11 and Piezo1 to integrate both mechanical and biochemical cues highlights the need for well-controlled *in vitro* experimental systems that can recapitulate mechanical and biochemical properties of tissue to better understand the relationships driving diseases like IPF.

Biomaterial-based *in vitro* models have become powerful platforms for modeling tissue microenvironments.^38–42^ Among these, hydrogels have been particularly valuable for studying cellular behavior within physiologically relevant settings, offering critical insights into how matrix mechanics regulate processes ranging from cell proliferation to stem cell fate.^41,43–49^ These water-swollen polymer networks can be fabricated from a range of materials, and the incorporation of orthogonal chemistries enables precise control and tunability of both static and dynamic mechanical properties. Independent control over properties such as material stiffness and viscoelasticity enables the decoupling of individual mechanical cues, revealing distinct and critical roles for each in regulating cell behavior. Studies from our lab and others have leveraged this tunability to demonstrate that stiffer substrates promote fibroblast activation as measured by increased cell spread areas, more irregular and spread morphologies, punctate focal adhesion formation, and expression of pro-fibrotic markers like type I collagen and alpha smooth muscle actin (LSMA) stress fibers.^16,47–53^ Control over substrate mechanics is particularly important for modeling fibrotic diseases such as IPF, where pathological progression involves changes in both the biochemical composition and mechanical properties of the ECM. By tuning these parameters in hydrogels, it becomes possible to dissect how specific mechanical features— either alone or in combination with biochemical signals—drive fibroblast activation, macrophage polarization, and other processes central to fibrogenesis.

We recently identified a compelling link between IL-6 signaling and M2 macrophage–mediated fibroblast activation, highlighting this pathway as a central driver of profibrotic communication^16^. Building on this, CDH11 and Piezo1 have emerged as two candidate regulators that may sustain and amplify this signaling axis.^33–37^ To probe their relative contributions, we employed a hyaluronic acid (HA)–based hydrogel system previously developed in our lab to mimic the mechanics of normal and fibrotic lung tissue.^47–49^ Using this platform, we investigated how CDH11 and Piezo1 influence fibroblast activation and IL-6–mediated macrophage–fibroblast crosstalk. This integrative approach begins to unravel the multifaceted interactions driving fibroblast activation in IPF and offers new insight into potential therapeutic targets.

## RESULTS AND DISCUSSION

### Piezo1 and Cadherin-11 expression increase with hydrogel stiffness and M2 macrophage co-culture

Given that Piezo1 and CDH11 not only mediate M2 macrophage-fibroblast crosstalk but are also linked to IL-6 signaling, a signaling pathway shown to drive pro-fibrotic fibroblast behavior, we first sought to examine their responses to physiologically relevant substrates and co-culture with M2 macrophages.^16,24,25,33–37^ Utilizing a hyaluronic acid (HA)-based hydrogel system previously developed in our lab,^47,48^ we fabricated soft (2 kPa) viscoelastic and stiff (25 kPa) elastic hydrogels to mimic normal and fibrotic lung, respectively. We seeded human lung fibroblasts on hydrogels with or without the addition of M2 macrophages to probe how substrate stiffness and macrophage co-culture influence Piezo1 and CDH11 expression (**Fig. 1**). We observed significantly increased cell area and more elongated, spindle-like morphology for fibroblasts seeded on 25 kPa elastic substrates and those cultured with M2 macrophages, regardless of substrate mechanics (**Fig. 1B, 1C**). These results support previous work from our lab, demonstrating that increased hydrogel stiffness activates fibroblasts and that M2 macrophage co-culture can override mechanical cues to promote fibroblast activation.^16^ Additionally, we observed a significant increase in Piezo1 (**Fig. 1D**) and CDH11 (**Fig. 1E**) expression in these groups, supporting their role in fibroblast activation related to both substrate mechanics and M2 macrophage signaling. Together, these results highlight the role of Piezo1 and CDH11 in mediating stiffness- and macrophage-driven fibroblast activation and underscore their potential roles in pro-fibrotic signaling.

**Figure 1:**
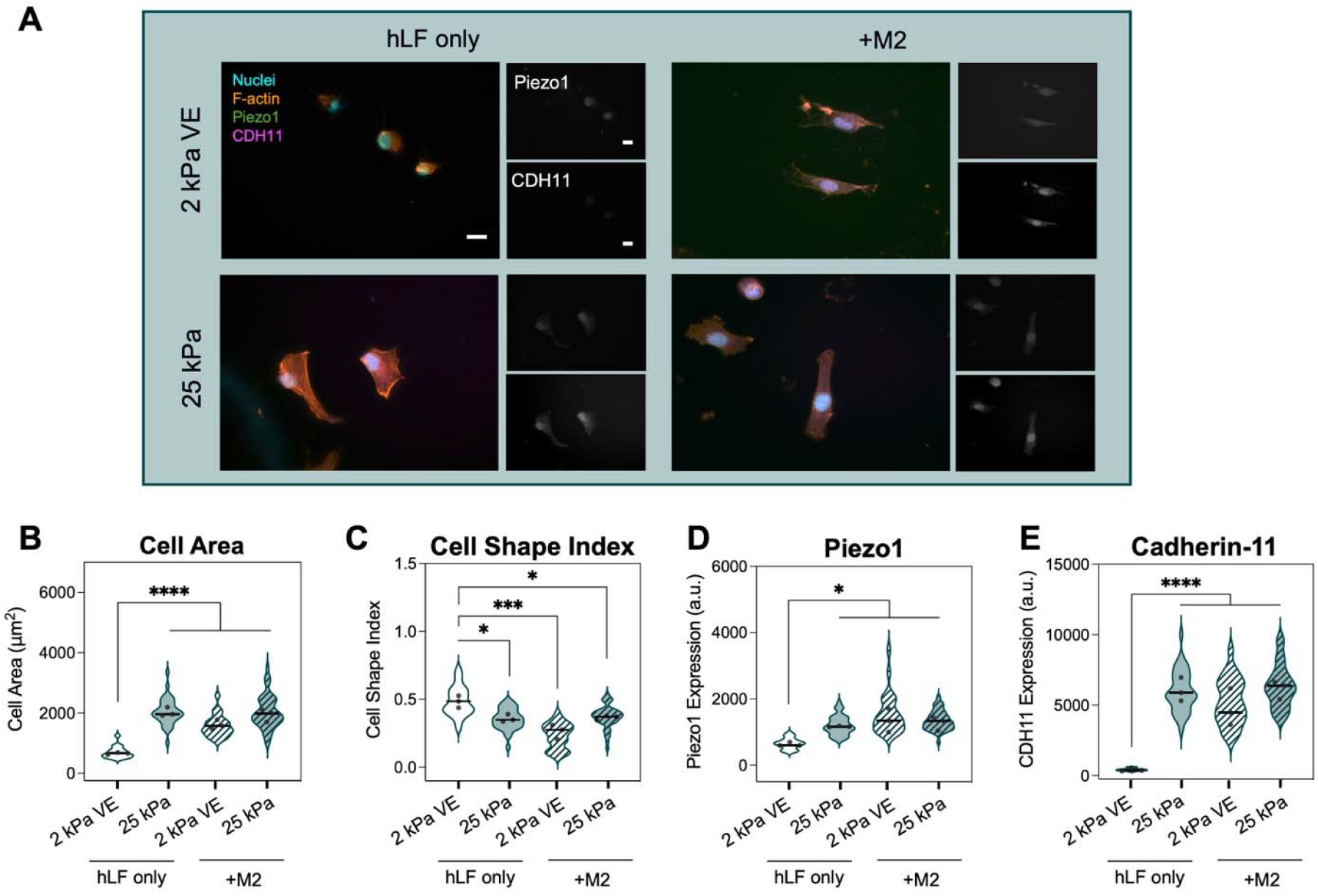
Piezo1 and CDH11 expression increase with hydrogel stiffness and M2 macrophage co-culture. A) Representative images of human lung fibroblasts (hLFs) seeded on 2 kPa viscoelastic (VE) and 25 kPa elastic substrates with or without the addition of M2 macrophages. Scale bars = 20 μm. B) Fibroblasts cultured on 25 kPa elastic hydrogels and with M2 macrophages exhibited significantly increased spread areas compared to those cultured alone on 2 kPa VE substrates. C) Fibroblasts on stiff hydrogels and in M2 co-culture adopted a more elongated, spindle-like morphology, reflected by significantly reduced cell shape index. D) Piezo1 expression significantly increases in fibroblasts cultured on 25 kPa hydrogels and those co-cultured with M2 macrophages. E) CDH11 expression significantly increases in fibroblasts cultured on 25 kPa hydrogels and those co-cultured with M2 macrophages. Each point represents one hydrogel average, *n* = 3 hydrogels per group, 45-104 individual cells per group. Statistical analyses performed via two-way ANOVA with Tukey’s HSD post hoc testing. \*\*\*\**p* < 0.0001, \*\*\**p* < 0.001, \**p* < 0.05.

### Blocking interleukin-6 (IL-6) signaling reduces cadherin-11, but not Piezo1, expression

IL-6 signaling has been established as a contributor to fibroblast activation, acting through a STAT3-dependent pathway similar to TGF-β1 signaling.^20,54,55^ In the context of IPF, inhibition of IL-6 has been shown to attenuate fibrosis progression in animal models of pulmonary fibrosis.^56,57^ Building on our recent finding that IL-6 is a key mediator of M2 macrophage-induced fibroblast activation,^16^ we next sought to investigate how IL-6 signaling intersects with mechanotransduction and cell-cell crosstalk. Specifically, we wanted to examine the effect of IL-6 inhibition on the expression of Piezo1 and CDH11.

To investigate this, wild type (WT) human lung fibroblasts and M2 macrophages were harvested from culture flasks, then either immediately seeded on 2 kPa viscoelastic and 25 kPa elastic hydrogels or incubated on ice with the IL-6R antibody, tocilizumab, to prevent IL-6 signaling. After 48 hours of culture, media was collected to quantify IL-6 via ELISA, and cells were stained and imaged to quantify cell spreading, morphology, and Piezo1 and CDH11 expression (**Fig. 2**). As observed previously, WT fibroblasts on stiff substrates and in co-culture with M2 macrophages exhibited increased activation, reflected by larger cell areas, more elongated morphology, and elevated CDH11 expression (**Fig. 2B–E**). In contrast, inhibition of IL-6 led to decreased fibroblast spread areas, promoted more rounded cell morphologies, and decreased expression of CDH11 in M2 co-cultures compared to control groups, consistent with prior work from our group.^16^ Notably, while Piezo1 expression showed a moderate increase in fibroblast-only cultures in response to IL-6 inhibition, its levels in M2 co-cultures were not significantly altered compared to untreated controls (**Fig. 2D**). These results suggest a connection between IL-6 signaling and CDH11 expression in fibroblast activation, while Piezo1 appears to operate more independently, indicating potentially distinct regulatory pathways.

**Figure 2:**
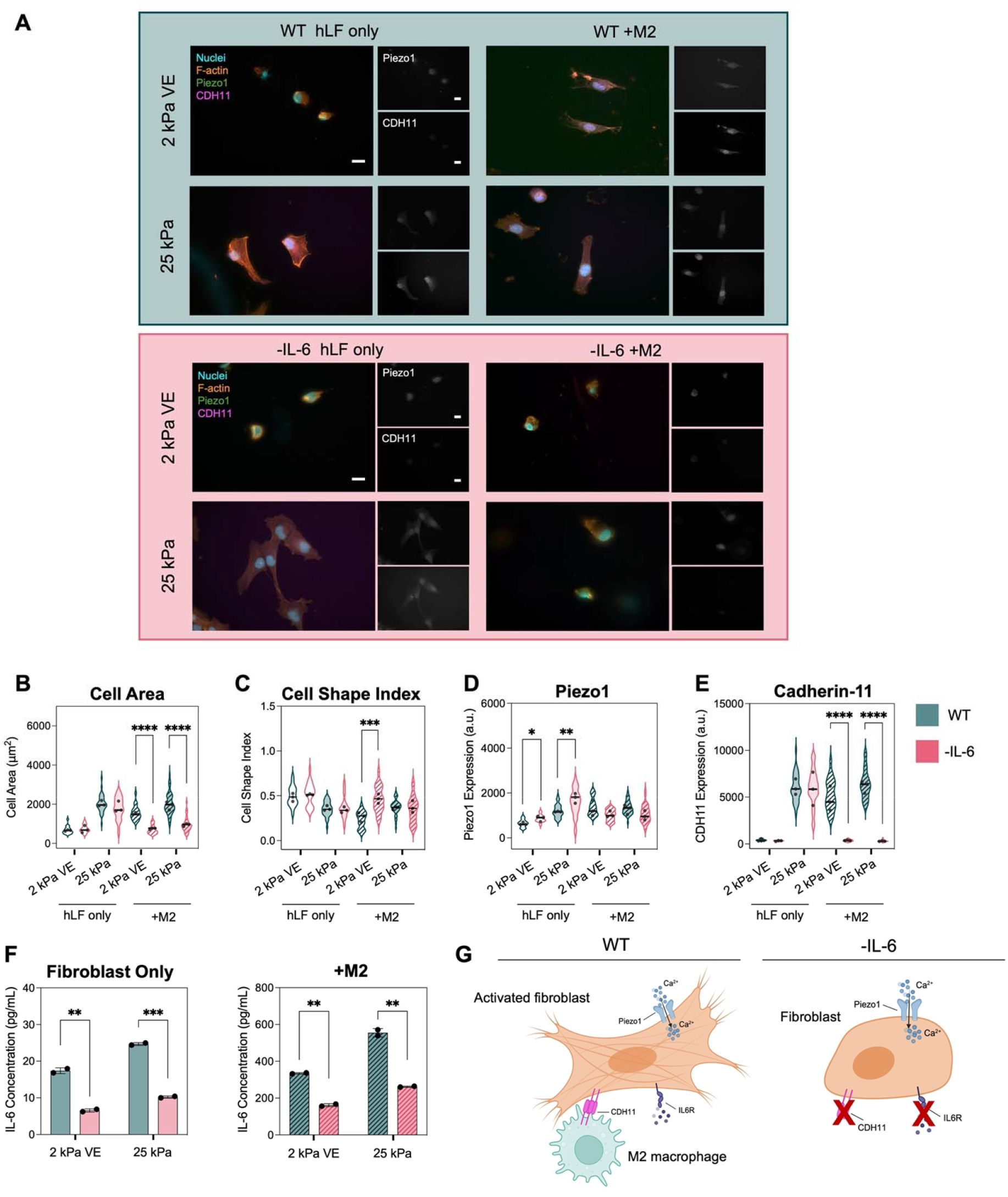
IL-6 signaling drives macrophage-mediated fibroblast activation and influences CDH11 expression. A) Representative images of wild type (WT) human lung fibroblasts (hLFs, top panel) and IL-6 inhibited (-IL-6) fibroblasts (bottom panel) seeded on 2 kPa viscoelastic (VE) and 25 kPa elastic substrates after 48 hours of culture. Scale bars = 20 μm. B) WT fibroblasts exhibited increased spread areas on 25 kPa hydrogels and in co-culture with M2 macrophages regardless of hydrogel stiffness. Conversely, IL-6 inhibited fibroblasts displayed reduced spreading in M2 macrophage co-culture groups, regardless of hydrogel stiffness. C) Cell circularity evaluation, measured by cell shape index, showed fibroblasts exhibited rounder morphologies in M2 macrophage co-cultures on 2 kPa VE gels when IL-6 was inhibited. D) WT fibroblasts showed increased expression of Piezo1 on 25 kPa substrates and in co-culture with M2 macrophages independent of hydrogel stiffness. IL-6 inhibited fibroblasts exhibited increased levels of Piezo1 expression in fibroblast only cultures compared to WT controls. E) WT fibroblasts showed higher levels of CDH11 expression on 25 kPa substrates and in M2 macrophage co-culture. IL-6 inhibited fibroblasts exhibited increased levels of CDH11 on 25 kPa hydrogels when cultured alone. However, IL-6 inhibited fibroblasts co-cultured with M2 macrophages displayed very little CDH11 expression, comparable with fibroblasts cultured alone on 2 kPa VE hydrogels. F) IL-6 inhibited fibroblast cultures exhibited decreased levels of IL-6 compared to WT controls, independent of stiffness or macrophage co-culture. G) We propose that inhibiting IL-6 signaling blocks M2 macrophage–induced fibroblast activation and lowers CDH11 expression, whereas Piezo1 signaling appears relatively unaffected. Each point represents one hydrogel average, *n* = 3 hydrogels per group, 47-104 individual cells per group. Statistical analyses performed via two-way ANOVA with Tukey’s HSD post hoc testing. \*\*\*\**p* < 0.0001, \*\*\**p* < 0.001, \*\**p* < 0.01, \**p* < 0.05.

### Piezo1 mediates early calcium-dependent responses in fibroblast activation

Since IL-6 signaling does not appear to significantly influence Piezo1 expression in lung fibroblasts, we next sought to investigate alternative pathways. Given that Piezo1 activation promotes calcium influx and that CDH11 function is calcium-dependent, we investigated the relative influence of Piezo1 and CDH11 signaling on the nuclear localization of the calcium-dependent transcription factor nuclear factor of activated T-cells 1 (NFAT1). NFAT1 was selected as a marker of calcium-mediated signaling as it is activated by intracellular calcium influx via calcineurin-dependent dephosphorylation.^62–64^ We utilized the stretch activated ion channel (SAC) inhibitor GsMTx4, a well-established inhibitor of Piezo1,^25,58–61^ to pharmacologically block Piezo1 activity in WT fibroblasts cultured on 2 kPa viscoelastic or 25 kPa elastic hydrogels, with or without M2 macrophages. WT fibroblasts cultured without inhibitor served as controls. Because there is no standardized small-molecule inhibitor of CDH11, we instead developed a CDH11 knockout (KO) fibroblast line to assess its role in fibroblast activation (see **Supplemental File 1**). After 4 hours, cells were fixed and stained to evaluate fibroblast spreading, morphology, and expression of CDH11 and NFAT1 (**Fig. 3**). At this early timepoint, NFAT1 nuclear localization was elevated in WT fibroblasts cultured on 25 kPa substrates and in co-culture with M2 macrophages (**Fig. 3A**). Piezo1 inhibition significantly reduced nuclear NFAT1 localization and cell area in these groups, while CDH11-KO fibroblasts displayed responses more similar to WT controls (**Fig. 3A, 3B**). These findings indicate that Piezo1 activity is linked to early NFAT1 activation, whereas CDH11 may not yet be engaged.

**Figure 3:**
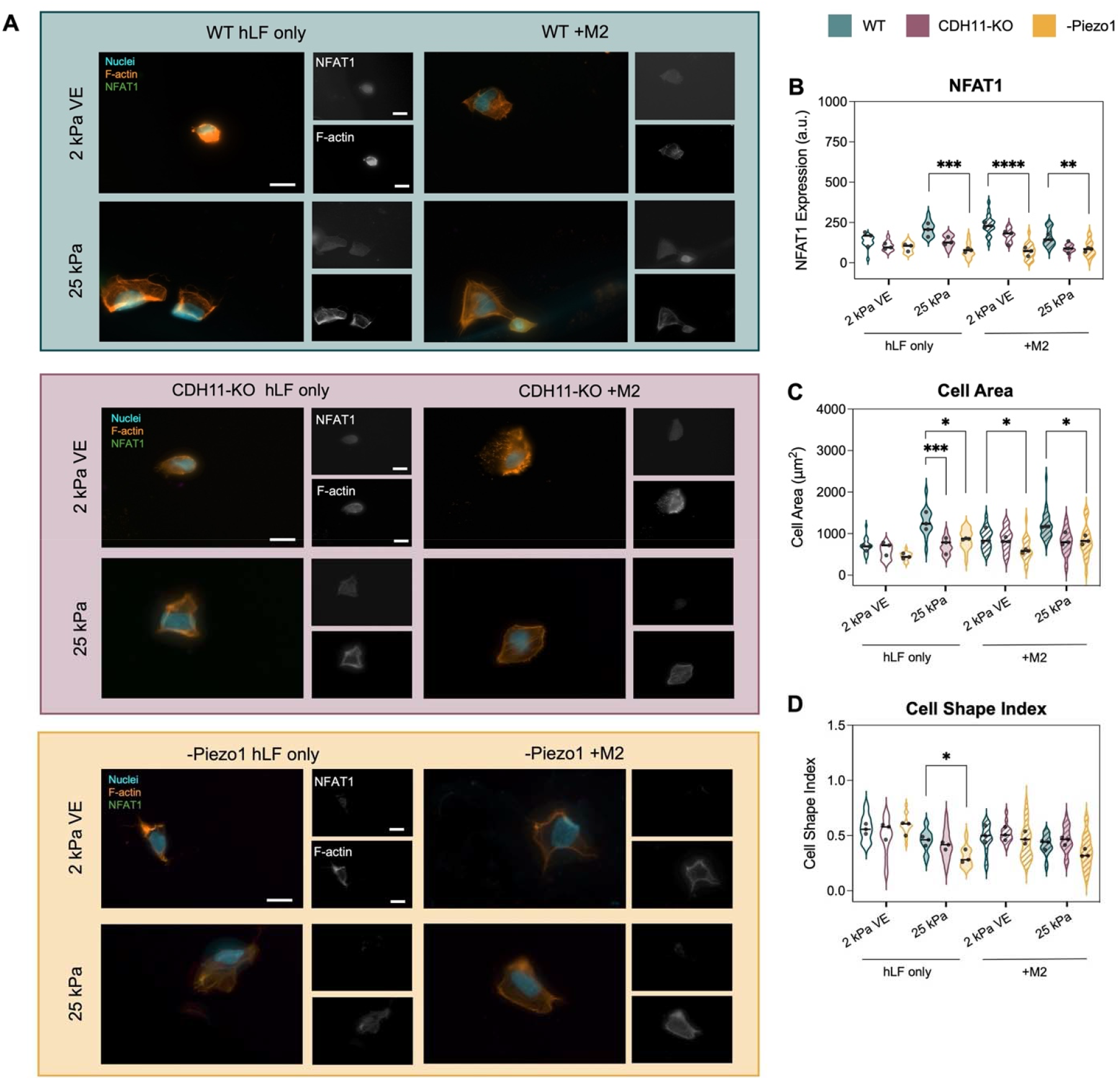
Piezo1 mediates calcium-dependent NFAT1 nuclear localization. A) Representative images of wild type (WT) human lung fibroblasts (hLFs, top panel), cadherin-11 knockout (CDH11-KO) fibroblasts (middle panel), and Piezo1-inhibited fibroblasts (bottom panel) seeded on 2 kPa viscoelastic (VE) and 25 kPa elastic substrates after 4 hours of culture. Scale bars = 20 μm. B) WT fibroblasts displayed higher levels of calcium-mediated nuclear NFAT1 expression when cultured on 25 kPa substrates and co-cultured with M2 macrophages. CDH11-KO fibroblasts did not have significantly different NFAT1 expression compared to WT controls in any culture group. Piezo1-inhibited cells exhibited significantly less nuclear NFAT1 expression in all groups compared to WT controls. C) WT fibroblasts exhibited increased spreading on 25 kPa hydrogels and in co-culture with M2 macrophages regardless of hydrogel stiffness compared to fibroblast only cultures on 2 kPa VE hydrogels. CDH11-KO cells followed the same trends as WT fibroblasts, however in fibroblast only cultures on 25 kPa hydrogels they did not spread to the same extent as controls. Piezo1-inhibited fibroblasts also followed a similar trend as WT controls; however, they did not spread to the same extent in any of the groups. D) All groups exhibited similar cell morphologies as measured by cell shape index. While Piezo1-inhibited fibroblasts exhibited slightly more elongated cell shapes compared to WT controls on 25 kPa hydrogels, this trend was not consistent across culture types. Each point represents one hydrogel average, *n* = 3 hydrogels per group, 37-59 individual cells per group. Statistical analyses performed via two-way ANOVA with Tukey’s HSD post hoc testing. \*\*\*\**p* < 0.0001, \*\*\**p* < 0.001, \*\**p* < 0.01, \**p* < 0.05.

Similar to 48-hour cultures, we observed increased fibroblast spreading on 25 kPa substrates and in M2 macrophage co-cultures in WT fibroblasts (**Fig. 3B**) after 4 hours. However, fibroblasts cultured on 2 kPa viscoelastic substrates with M2 macrophages did not spread to the same extent as those on 25 kPa hydrogels, in contrast to the 48-hour results (**Fig. 1B, 3C**). This suggests that fibroblast activation driven by M2 macrophage co-culture on soft substrates is a progressive process that requires extended signaling dynamics beyond this 4-hour timepoint. Consistent with this, no significant changes in cell morphology were observed in WT fibroblasts across hydrogel stiffness or co-culture conditions at this early time point (**Fig. 3D**), indicating that cells are still undergoing progressive activation.

Furthermore, Piezo1-inhibited groups supported decreased cell areas and reduction in nuclear NFAT1 staining, while CDH11-KO cells are more similar to WT controls (**Fig. 3B, 3C**). Notably, compared to 48-hour time points, CDH11 expression is significantly lower after 4 hours of culture, while Piezo1 levels are comparable across timepoints (**Fig. S1**). These findings support the idea that CDH11 engagement does not occur within the first 4 hours of culture, consistent with previous studies showing a lack of CDH11 expression in ‘naïve’ fibroblasts and its increased expression in more mature, contractile myofibroblasts.^25,65^ Overall, these results suggest that Piezo1-dependent nuclear localization of the calcium-dependent transcription factor NFAT1 may play an important role in early fibroblast responses to substrate stiffness and M2 macrophage co-culture, while CDH11 could act as a downstream effector that is later engaged to support more sustained or mature fibroblast activation.

### Piezo1 inhibition moderately reduces cadherin-11 expression but does not significantly alter fibroblast activation

Given the well-established role of Piezo1 in cellular mechanotransduction,^66,67^ the increased levels of Piezo1 expression we quantified in activated fibroblasts, and the role of Piezo1 we observed in mediating early calcium-dependent NFAT1 signaling, we next wanted to investigate how fibroblast activation was influenced by Piezo1 inhibition. We again seeded WT fibroblasts on 2 kPa viscoelastic and 25 kPa elastic hydrogels, with or without the addition of M2 macrophages. After allowing cells to adhere overnight, media was changed to either fresh media or media supplemented with 2 mM GsMTx4 to inhibit Piezo1 activity for an additional 24 hours. After this time, we measured IL-6 levels in culture media via ELISA and quantified fibroblast activation by measuring cell spread area, shape morphology, and expression of type I collagen, Piezo1, and CDH11 in WT and Piezo1-inhibited cells.

We found that WT fibroblasts maintained consistent trends of activation on 25 kPa elastic substrates and in the presence of M2 macrophages independent of stiffness, supporting previous results and our earlier experiments in this manuscript (**Fig. 4, S2**). Piezo1 inhibition did not significantly influence fibroblast spread area, shape, or type I collagen expression compared to the WT controls, suggesting that Piezo1 blockade alone is insufficient to suppress fibroblast activation (**Fig. 4, S2**). While Piezo1 inhibition led to a slight increase in IL-6 secretion, this did not translate into functional differences, as we did not observe corresponding increases in fibroblast activation markers (**Fig. 4F**). This modest rise in IL-6 may reflect crosstalk with MAPK signaling, a pathway regulated by both Piezo1 and IL-6 activity, where perturbation of one input could modulate the other.^37,68^ Taken together, these findings suggest that while Piezo1 influences early calcium-mediated NFAT1 signaling, it is not essential for fibroblast activation under these conditions.

**Figure 4:**
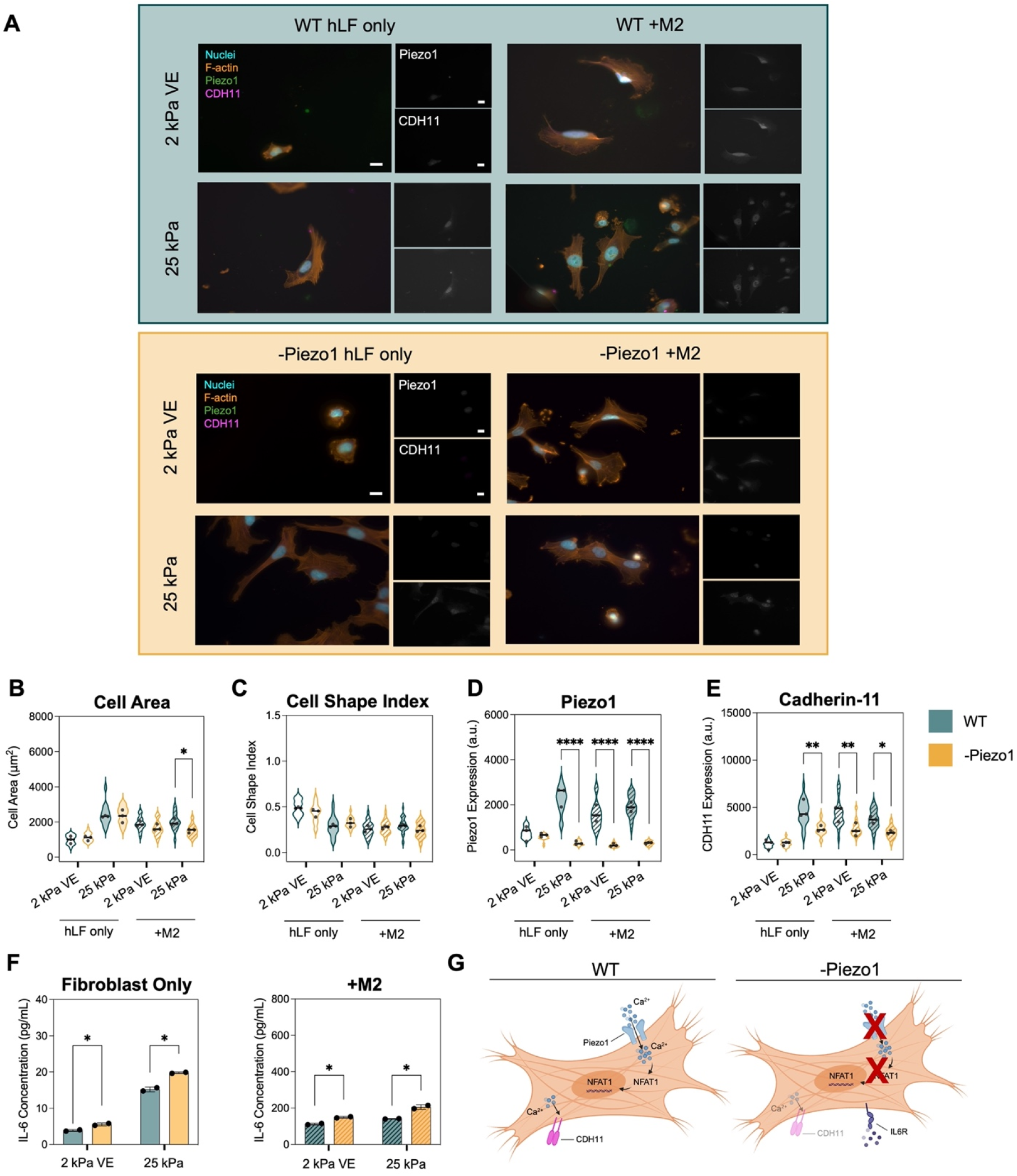
Piezo1 inhibition does not significantly alter stiffness- or macrophage-induced fibroblast activation. A) Representative images of wild type (WT) fibroblasts (top panel) and Piezo1 inhibited (-Piezo1) fibroblasts (bottom panel) seeded on 2 kPa viscoelastic (VE) and 25 kPa elastic substrates after 48 hours of culture. Scale bars = 20 μm. B) WT and Piezo1 inhibited fibroblasts exhibited increased spread areas on 25 kPa hydrogels and in co-culture with M2 macrophages regardless of hydrogel stiffness. Though there was a slight decrease in Piezo1-inhibited fibroblast spreading in co-culture with M2 macrophages on 25 kPa hydrogels, these cells still showed significantly higher areas than those cultured alone on 2 kPa substrates. C) Cell circularity evaluation, as measured by cell shape index, showed no significant difference in cell morphologies between Piezo1-inhibited fibroblasts and WT controls. D) WT fibroblasts displayed increased Piezo1 expression on 25 kPa substrates and in co-culture with M2 macrophages independent of hydrogel stiffness. Piezo1-inhibited fibroblasts had very little Piezo1 expression in any group, indicating successful inhibition with GsMTx4. E) WT fibroblasts had higher levels of CDH11 expression on 25 kPa substrates and in M2 macrophage co-culture. Piezo1-inhibited fibroblasts showed moderate reductions in overall CDH11 expression levels. F) Piezo1-inhibited fibroblast cultures exhibited marginally increased IL-6 levels compared to WT controls. G) We propose that Piezo1 inhibition prevents calcium influx and downstream NFAT1 signaling, leading to moderate reductions in CDH11 expression, but does not prevent fibroblast activation. Each point represents one hydrogel average, *n* = 3 hydrogels per group, 51-95 individual cells per group. Statistical analyses performed via two-way ANOVA with Tukey’s HSD post hoc testing. \*\*\*\**p* < 0.0001, \*\**p* < 0.01, \**p* < 0.05.

Notably, Piezo1 inhibition led to moderate reduction in CDH11 expression, suggesting a potential mechanistic relationship between Piezo1 activity and CDH11 regulation (**Fig. 4E**). Since Piezo1 drives early calcium signaling that precedes CDH11 expression, and CDH11 activation is calcium-dependent, it is likely that reduced calcium influx contributes to the decrease in CDH11 observed in Piezo1-inhibited fibroblasts. Furthermore, the progressive increase in fibroblast activation from 4 to 48 hours, together with the significant increase in CDH11 expression at 48 hours (**Fig. S1**), indicates that CDH11 may play a more central role than Piezo1 in sustaining fibroblast activation.

### Cadherin-11 KO significantly reduces Piezo1 expression, interleukin-6 secretion, and fibroblast activation

Given our observations that CDH11 expression increases with progressive fibroblast activation and is IL-6-dependent, we next wanted to investigate changes in fibroblast behavior upon knocking out CDH11. WT and CDH11-KO fibroblasts were seeded on 2 kPa viscoelastic and 25 kPa elastic hydrogels, with or without the addition of M2 macrophages. Media samples were collected to measure IL-6 concentration and fibroblast activation was assessed by measuring spread area, morphology, and expression of type I collagen, Piezo1, and CDH11. Consistent with previous findings, we observed increased fibroblast spread area, more elongated and spindle-like morphology, and increased expression of type I collagen, Piezo1, and CDH11 in WT fibroblasts seeded on stiffer elastic substrates and in M2 macrophage co-cultures (**Fig. 5, S3**). Notably, we found that CDH11-KO fibroblasts exhibited decreased spreading, rounder cell morphologies, and decreased levels of type I collagen and Piezo1, regardless of stiffness or macrophage co-culture (**Fig. 5, S3**). CDH11-KO returns levels of these activation markers to those observed in WT fibroblasts seeded on 2 kPa viscoelastic hydrogels, suggesting that loss of CDH11 is sufficient to prevent fibroblast activation in the presence of mechanical and biochemical stimuli.

**Figure 5:**
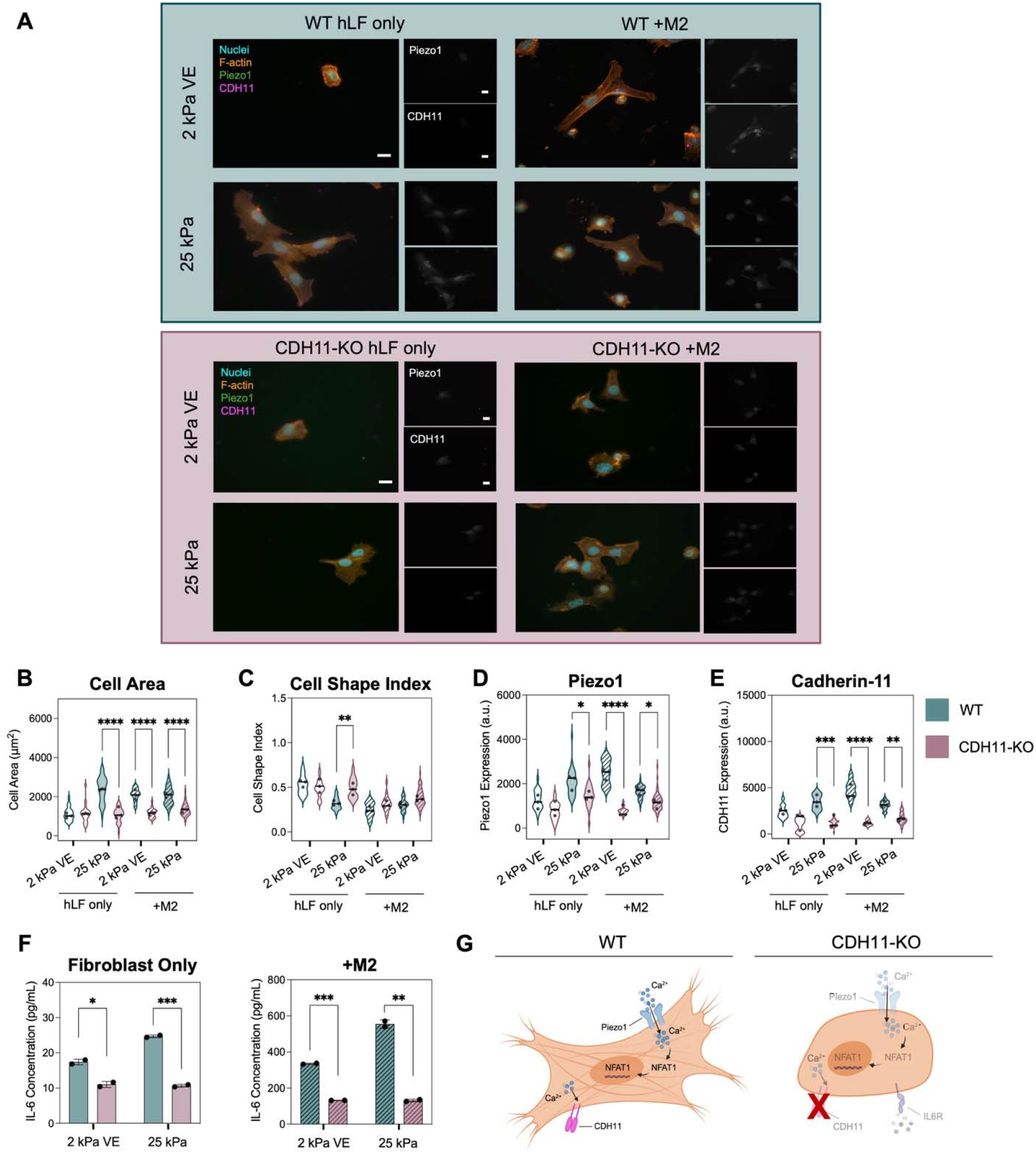
Cadherin-11 knockdown prevents stiffness- and M2 macrophage-dependent fibroblast activation. A) Representative images of wild type (WT) fibroblasts (top panel) and cadherin-11-knockout (CDH11-KO) fibroblasts (bottom panel) seeded on 2 kPa viscoelastic (VE) and 25 kPa elastic substrates after 48 hours of culture. Scale bars = 20 μm. B) WT fibroblasts exhibited increased spread areas on 25 kPa hydrogels and in co-culture with M2 macrophages regardless of hydrogel stiffness. However, CDH11-KO fibroblasts showed significantly smaller spread areas in these conditions, indicating a lack of response to mechanical or macrophage-derived cues. C) Cell circularity evaluation, as measured by cell shape index, showed that CDH11-KO fibroblasts exhibited more rounded cell morphologies compared to WT controls. D) WT fibroblasts exhibited increased Piezo1 expression on 25 kPa substrates and in co-culture with M2 macrophages independent of hydrogel stiffness. CDH11-KO fibroblasts showed a significant reduction in Piezo1 expression in these same groups. E) WT fibroblasts displayed higher levels of CDH11 expression on 25 kPa substrates and in M2 macrophage co-culture, while CDH11-KO cells did not exhibit CDH11 expression as expected. F) CDH11-KO fibroblast cultures had decreased levels of IL-6 compared to WT controls, regardless of stiffness or macrophage co-culture, highlighting the reciprocal relationship between IL-6 and CDH11 engagement. G) We propose that CDH11-KO prevents both stiffness- and M2 macrophage-mediated fibroblast activation through inhibitory effects on Piezo1 and IL-6 signaling. Each point represents one hydrogel average, *n* = 3 hydrogels per group, 52-115 individual cells per group. Statistical analyses performed via two-way ANOVA with Tukey’s HSD post hoc testing. \*\*\*\**p* < 0.0001, \*\*\**p* < 0.001, \*\**p* < 0.01, \**p* < 0.05.

These results are consistent with studies from other groups that demonstrate CDH11 knockdown leads to reduced fibrosis and inflammation in *in vivo* models of pulmonary fibrosis, myocardial infarction, and rheumatoid arthritis.^23,33,34^ Additionally, while Piezo1 and CDH11 have both been implicated in fibrosis and macrophage crosstalk, there have not been studies exploring the relationship between these mediators.^22,24,25,27^ The reduction in Piezo1 we observed in response to CDH11 knockdown has not been previously reported but is not necessarily surprising, as CDH11 is the only cadherin known to mediate both cell-cell and cell-ECM crosstalk.^28,29^ Additionally, both CDH11 and Piezo1 have been implicated in the regulation of MAPK signaling pathways.^34,37,69,70^ Within this context, it is possible that CDH11 knockdown disrupts cell-ECM interactions and/or signaling pathways that also influence the engagement of Piezo1. Overall, these results emphasize the important role CDH11 plays in driving fibroblast activation and the influence it may have over other mechanotransduction pathways crucial to fibrogenesis.

Moreover, media from CDH11-KO cell cultures contained significantly lower levels of IL-6 compared to WT controls, further supporting the dynamic relationship between CDH11 activation and IL-6 signaling (**Fig. 5F**). This result, in addition to the decreased fibroblast activation observed in CDH11-KO cells and significant decrease in CDH11 expression in IL-6 inhibited fibroblasts (**Fig. 2E**), supports work from other groups that have shown that CDH11 engagement and IL-6 signaling have reciprocal relationships in cardiac^33^ and synovial^34^ fibroblasts. Combined with our findings that Piezo1 mediated-calcium signaling precedes CDH11 expression and that blocking Piezo1 or CDH11 activity leads to a reduction in the other’s expression, these results point to a novel mechanoinflammatory circuit between Piezo1, CDH11, and IL-6.

## CONCLUSIONS

Overall, this work demonstrates the interconnected relationships between Piezo1, IL-6, and CDH11 in driving fibroblast activation and identifies CDH11 as a key mediator of sustained activation. We demonstrate that both Piezo1 and CDH11 are upregulated in activated fibroblasts (**Fig. 1**) but found that only CDH11 expression is dependent on IL-6 signaling (**Fig. 2, 6A**), suggesting that Piezo1 contributes to fibroblast activation through a separate pathway. Given that Piezo1 functions as a mechanosensitive calcium channel, we hypothesized that it may contribute to early calcium-dependent signaling events that influence fibroblast activation. Indeed, at 4-hour time points, Piezo1 influenced calcium-mediated signaling as measured by nuclear NFAT1 levels (**Fig. 3**). Additionally, we found that CDH11 expression was significantly reduced at 4-hours compared to 48-hours of culture, while Piezo1 levels were consistent over this timeframe (**Fig. S1**). Given that CDH11 activation is calcium-dependent, we propose that Piezo1 engagement contributes to CDH11 upregulation at 48-hour timepoints. Consistent with this, we found that Piezo1 inhibition did not prevent fibroblast activation, however its inhibition results in a decrease of CDH11 expression (**Fig. 4, 6B**).

**Figure 6:**
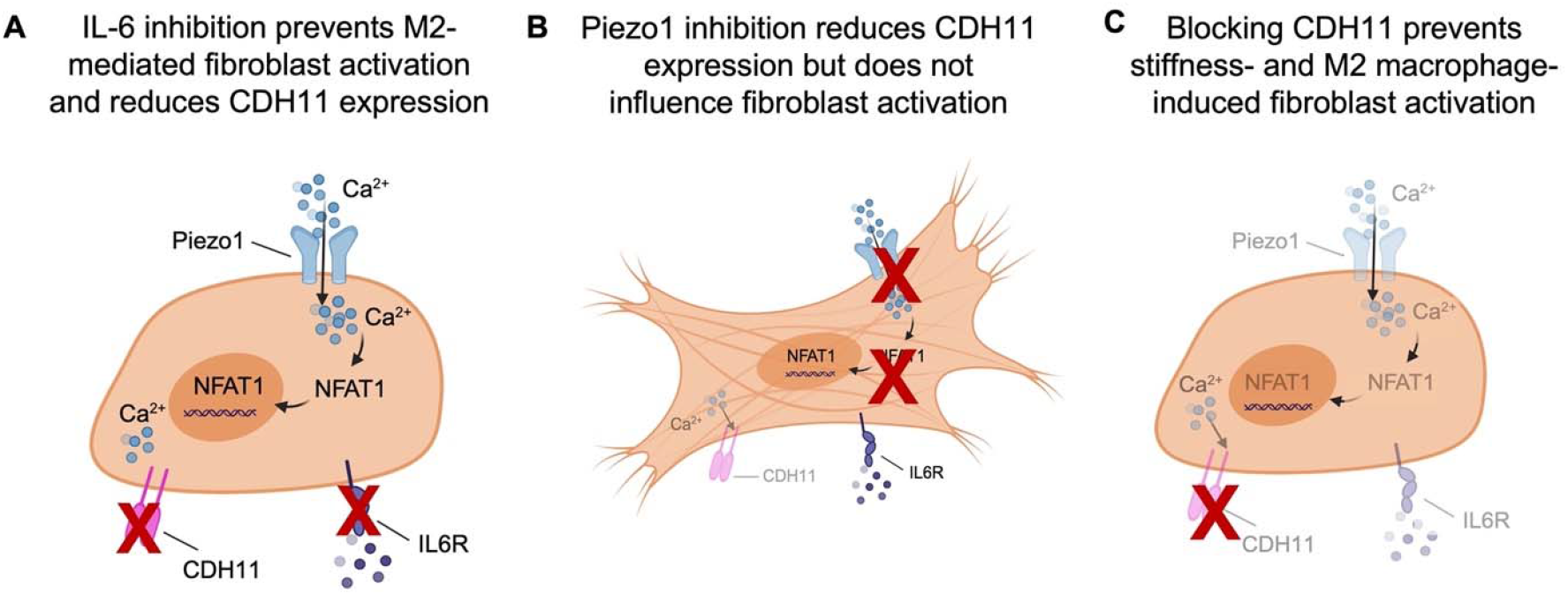
Summary of relationships between Piezo1, IL-6 signaling, and CDH11. A) Inhibiting IL-6 signaling prevents M2 macrophage-mediated fibroblast activation, reducing cell spreading and CDH11 expression. B) Piezo1 inhibition moderately reduces CDH11 expression, likely due to a reduction in calcium influx, but does not prevent fibroblast activation. C) CDH11 knockdown prevents stiffness- and M2 macrophage-mediated fibroblast activation, highlighting its role in sustaining profibrotic phenotypes.

In contrast, we found that loss of CDH11 has a more pronounced effect on fibroblast activation. CDH11 knockdown prevented both stiffness- and M2 macrophage-induced fibroblast activation, reduced Piezo1 expression, and resulted in decreased levels of IL-6 secretion (**Fig. 5, 6C**). Moreover, CDH11 expression increases in parallel with fibroblast activation between 4 and 48 hours of culture, further supporting its role in amplifying or sustaining fibroblast activation (**Fig. S1**). Together, these findings establish CDH11 as a central mediator that integrates mechanical and inflammatory cues, highlighting its significant influence over both mechanosensitive channels and cytokine signaling.

In summary, our results uncover a dynamic interplay between Piezo1, IL-6, and CDH11, revealing a previously unrecognized signaling axis that drives fibroblast activation in pulmonary fibrosis. Within this axis, CDH11 emerges as the dominant mediator of sustained activation, working in concert with IL-6, while Piezo1 plays a more supportive role in initiating calcium-dependent signaling.

## ACKNOWLEDGMENTS

We thank Lily Burns and Ethan Yu for their assistance maintaining cells and synthesizing hydrogel materials used in this work, as well as Dr. Dan Abebabyehu and Caliari Lab members for helpful discussions, support, and writing feedback. This work was supported by the NIH (R35GM138187) and NSF (CAREER DMR/BMAT 2046592, GRFP to L.R.A.). Flow cytometry data for this manuscript were generated in the University of Virginia Flow Cytometry Core Facility (RRid:SCR_017829) that is partially supported by the NCI Grant P30-CA044579. Cell sorting was performed on the Cytek Aurora CS funded through the NIH S10 instrumentation grant (1S10OD034355-01). The content is solely the responsibility of the authors and does not necessarily represent the official views of the National Institutes of Health.

## EXPERIMENTAL SECTION

### Norbornene-modified HA (NorHA) synthesis

NorHA was synthesized according to previously described methods.^71^ Briefly, sodium hyaluronate (Lifecore, 82 kDa) was first converted to a tetrabutylammonium salt (HA-TBA) using Dowex 50W proton-exchange resin. The exchange proceeded for 2 h before filtering out the resin and adjusting the pH to 7.05 using TBA-OH. The product was then frozen, lyophilized, and analyzed using ^1^H NMR (Bruker Neo 400 MHz NMR Spectrometer) (**Fig. S4**). HA-TBA was dissolved in anhydrous dimethyl sulfoxide (DMSO) and reacted with 5-norbornene-2-methylamine, coupling to the carboxylic acid of HA with benzotriazole-1-yl-oxy-tris- (dimethylamino)-phosphonium hexafluorophosphate (BOP). The reaction proceeded for 2 h at 25°C before quenching with cold reverse osmosis (RO) water. The product was dialyzed (molecular weight cutoff 6–8 kDa) in brine for 5 days, filtered to remove side products, dialyzed against brine an additional 5 days, dialyzed against RO water for 2 days, frozen, and lyophilized. The degree of modification was determined to be 29% by ^1^H NMR (**Fig. S5**).

### β-CD-HDA synthesis

β-CD hexamethylene diamine (β-CD-HDA) was synthesized using previously reported methods.^72^ Briefly, β-CD was dissolved in RO water and degassed, *p*-toluenesulfonyl chloride (TosCl) dissolved in acetonitrile was then added dropwise to the β-CD solution (5:4 molar ratio of TosCl:CD) and proceeded to react for 2 h on ice. NaOH solution was added dropwise to the reaction (3.1:1 molar ratio of NaOH to CD), then proceeded for 30 min at 25°C. Ammonium chloride was then added until the reaction was at a pH of 8.5, the solution was again cooled on ice before precipitation using cold water and acetone, then dried overnight. The CD-Tos product was charged with excess hexamethylene diamine (HDA) (4 g/g CD-Tos) and dissolved in dimethylformamide (DMF) (5 mL/g CD-Tos). The reaction proceeded under nitrogen for 12 h at 80°C, then was precipitated in cold acetone (5 × 50 mL acetone/1 g CD-Tos), washed with cold diethyl ether (3 × 100 mL), and dried. The β-CD-HDA product was confirmed using ^1^H NMR (**Fig. S6**).

### CDHA synthesis

To synthesize β-CDHA, β-CD-HDA was coupled to HA-TBA using BOP according to previously described methods. The β-CD-HDA and HA-TBA were dissolved in anhydrous DMSO, followed by dropwise addition of BOP. The reaction proceeded for 3 h at 25°C before quenching with cold RO water. The solution was then dialyzed against brine (molecular weight cutoff 6–8 kDa) for 5 days, filtered to remove side products, dialyzed against brine for an additional 5 days, then dialyzed against RO water for 2 days, frozen, and lyophilized. The degree of modification was determined to be 25% by ^1^H NMR (**Fig. S7**).

### HA hydrogel fabrication and mechanical characterization

Glass coverslips (15 mm diameter) were thiolated to covalently bond norbornene groups within hydrogel precursor solutions using (3-mercaptopropyl) trimethoxysilane. Hydrogels were crosslinked between the thiolated and an untreated (12 mm diameter) coverslip. Prior to hydrogel precursor preparation, thiolated adamantane peptide (Ad-KKKCG) and dithiothreitol (DTT) crosslinker were reduced using tris(2-carboxyethyl)phosphine (TCEP) at a 0.5:1 molar ratio. To prepare 3 wt% HA soft viscoelastic (2.5 kPa) hydrogels, CDHA and the thiolated adamantane peptide were first combined (1.2:1 Ad:CD molar ratio) to enable guest–host complexation. NorHA, DTT (0.1 thiol:norbornene ratio), a thiolated RGD peptide (GCGYGR□G□D□SPG, 1 mM, GenScript), lithium acylphosphinate photoinitiator (LAP, 1 mM), and phosphate-buffered saline (PBS) (to reach a final volume of 17.5 μL) were subsequently added. Stiff elastic (25 kPa) 5 wt% HA hydrogels were fabricated similarly, without the inclusion of CDHA or the thiolated adamantane peptide and with increased DTT (0.4 thiol:norbornene ratio). After vortexing precursor solutions, hydrogels were photocrosslinked by exposing precursors to 365 nm UV light (5 mW cm^−2^) for 2 min using a VWR UV Crosslinker. After crosslinking, hydrogels were submerged in PBS and swelled overnight at 37°C before mechanical characterization or cell seeding.

Hydrogel mechanical characterization was performed via nanoindentation using an Optics11 Life Piuma nanoindenter (**Fig. S8**). A borosilicate glass probe with a diameter of 50 μm with a spring constant of 0.47 N/m was utilized for all tests. Indentations were made to a depth of 4 μm using a constant time ramp of 2 s. Storage (E′) and loss (E″) moduli were determined using dynamic mechanical analysis (DMA) at a frequency of 1 Hz at a depth of 4 μm at 3 points per hydrogel replicate.

### Cadherin-11 knock-out (CDH11-KO) cell line generation and characterization

Human lung fibroblasts (hTERT T1015 cell line purchased from Applied Biological Materials) were maintained in Dulbecco’s modified Eagle medium (DMEM) supplemented with 10 v/v% fetal bovine serum (FBS, Gibco) and 1 v/v% penicillin/streptomycin/amphotericin B (100 U mL^−1^ 100 μg mL^−1^ and 0.25 μg mL^−1^ final concentrations, respectively, Gibco) prior to transfection in 6-well tissue culture plates. VectorBuilder was utilized to generate an mCherry-tagged sgRNA/Cas9 vector targeting the CDH11 gene in human cells and provided corresponding DNA. Lipofectamine 3000 reagent (ThermoFisher, L3000001) was diluted in Opti-MEM medium (ThermoFisher, 31985070) and used to transfect fibroblasts according to manufacturer protocol. 5 ug/μL of DNA was added to each well, media was refreshed after 24 h to remove lipofectamine, and after 3 days cells were harvested for cell sorting via the University of Virginia Flow Cytometry core facility (**Supplemental File 1**).

### Cell culture

THP-1 human monocytic cells (TIB-202 cell line purchased from the American Type Culture Collection) were used before passage 7 for all experiments. Roswell Park Memorial Institute (RPMI) 1640 (Gibco) medium supplemented with 10 v/v% heat-inactivated FBS (Gibco), 1 v/v% penicillin/streptomycin/amphotericin B (100 U mL^−1^, 100 μg mL^−1^, and 0.25 μg mL^−1^ final concentrations, respectively, Gibco), 10 mM HEPES, 1 mM sodium pyruvate, 2.5 g L^−1^ D-glucose, and 0.05 mM β-mercaptoethanol was used for cell culture maintenance. Monocytes were differentiated to M0 macrophages by adding 50 ng mL^−1^ of phorbol 12-myristate 13-acetate (PMA) to fresh media and culturing for 24 h. Macrophages were then polarized to an M2 phenotype through the addition of IL-4 (40 ng mL^−1^; PeproTech) and IL-13 (20 ng mL^−1^; PeproTech) to fresh media. After 48 h, M2 macrophages were seeded onto hydrogels at a concentration of 3.0 x 10^5^ cells/hydrogel with either wild type human lung fibroblasts or CDH11 KO fibroblasts. Prior to seeding cells on hydrogels, hydrogels were sterilized for at least 2 h by germicidal UV irradiation and incubated in culture medium for 30 min.

Wild type human lung fibroblasts (hTERT T1015 cell line purchased from Applied Biological Materials) or CDH11-KO fibroblasts generated from the same line were utilized before passage 5 for all experiments. DMEM supplemented with 10 v/v% fetal bovine serum (FBS, Gibco) and 1 v/v% penicillin/streptomycin/amphotericin B (100 U mL^−1^ 100 μg mL^−1^ and 0.25 μg mL^−1^ final concentrations, respectively, Gibco) was used to maintain fibroblast culture. Fibroblasts were seeded on hydrogels at a concentration of 1.0 x 10^4^ cells per hydrogel and cultured with or without macrophages for 4 h or 48 h before fixing and immunocytochemistry.

### Immunocytochemistry, imaging, and analysis

Cells seeded on hydrogels were fixed using 10% neutral-buffered formalin for 15 min, followed by permeabilization using 0.1% Triton X-100 in PBS for 10 min. Background staining was blocked through incubation with bovine serum albumin (BSA; 3 w/v%) for 2 h at room temperature with gentle shaking. Primary antibodies against CDH11 (mouse monoclonal OB-cadherin/CDH11 antibody 16A, 1:200, ThermoFisher MUB0306P) and one of the following: Piezo1 (rabbit polyclonal Piezo1 antibody, 1:200, ThermoFisher 15939-1-AP), type 1 collagen (rabbit monoclonal anti-collagen I antibody, 1:200, Abcam ab138492), or NFAT1 (rabbit polyclonal NFAT1 antibody, 1:200, Cell Signaling Technologies 4389) were incubated with hydrogels overnight at 4°C. The next day, hydrogels were washed 3 times with PBS then incubated with rhodamine phalloidin (1:400, ThermoFisher R415) and secondary antibodies (AlexaFluor 488 goat anti-rabbit IgG, 1:400; AlexaFluor 647 goat anti-mouse, 1:400) for 2 h at room temperature protected from light. Hydrogels were again rinsed with PBS 3 times then stained with 4′,6-diamidino-2-phenylindole (DAPI, 1:5000, ThermoFisher D1306) for 5 min. After rinsing 3 times with PBS, hydrogels were stored protected from light at 4°C until imaging on a Zeiss AxioObserver 7 inverted microscope.

Cell area, cell shape index (form factor), and staining intensity of CDH11 and Piezo1 were evaluated using a CellProfiler (Broad Institute, Harvard/MIT) pipeline. Cell shape index measures cell circularity, where a perfect line and circle are assigned values of 0 and 1, respectively. This cell characteristic is quantified using the formula:

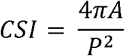

Nuclei were identified from a single channel DAPI image using adaptive thresholding, then overlaid with cells identified from a corresponding F-actin image to separate cells from background. Cells identified with equal nuclear and cellular area were assumed to be debris and removed before downstream analysis.

### Inhibitor experiments

Human lung fibroblasts were cultured as described above with or without the addition of M2 macrophages on 2.5 kPa viscoelastic or 25 kPa elastic hydrogels. Hydrogels were seeded as described above with 1.0 x 10^4^ fibroblasts and 3.0 x 10^5^ macrophages per hydrogel. Cells treated with the IL-6R antibody tocilizumab (abcam ab275982) were incubated with the antibody on ice in 2 v/v% FBS in PBS for 45 min, then washed 3 times with 2 v/v% FBS in PBS, before being resuspended in supplemented DMEM or RPMI media used for typical culture maintenance described above and seeded on hydrogels. Cells treated with the stretch-activated cation channel inhibitor GsMTx4 (R&D Systems 4912) were seeded and incubated overnight before switching to inhibitor containing media (2 mM GsMTx4) or fresh media for control wells. All groups were then cultured for an additional 4 h or 24 h before fixing and subsequent staining as described above.

### Quantification of IL-6 levels

After 48 h of culture, media samples were taken from wild type, CDH11 KO, Piezo1 inhibited, and IL-6 inhibited groups for both fibroblast only and M2 macrophage co-culture groups on 2.5 kPa viscoelastic and 25 kPa elastic hydrogels. Media samples were centrifuged to remove any cell debris before supernatants were flash frozen in liquid nitrogen and stored at −70°C until IL-6 quantification using a human IL-6 ELISA Kit (R&D Systems, D6050B). All samples and standards were run in duplicate according to manufacturer’s instructions and fluorescence intensity measured at 450 and 570 nm with a Tecan M200 plate reader.

### Statistical analysis

All statistical analyses were performed using GraphPad Prism. Outlier removal was performed using ROUT (Q = 1%) before downstream analysis. Two-way ANOVA with Tukey’s HSD post hoc analysis was used for comparisons between experimental groups. Experiments consisted of at least 3 hydrogels per group and/or a minimum of 30 individual cells quantified per group. Statistically significant differences are indicated by ^*, **, ***^, and ^****^ corresponding to *p* < 0.05, 0.01, 0.001, or 0.0001 respectively. Additional information regarding sample size or statistical analyses can be found in figure captions.

## Resource Availability

### Lead Contact

Further information and requests for resources and reagents should be directed to and will be fulfilled by the lead contact, Steven Caliari.

### Materials Availability

Hydrogel materials (NorHA, CDHA, etc.) in this study will be made available on request, but we may require a payment and/or a completed materials transfer agreement if there is potential for commercial application.

### Data and Code Availability

All data needed for the conclusions in the paper are present in the paper and the supplemental information. Additional data for the paper are available upon reasonable request from the lead contact.

## Author Contributions

Conceptualization, L.R.A. and S.R.C.; Methodology, L.R.A. and S.R.C.; Investigation, L.R.A.; Writing – Original Draft, L.R.A.; Writing – Review & Editing, L.R.A. and S.R.C.; Funding Acquisition, L.R.A. and S.R.C.

