## Supplemental Information for "Cadherin-11 integrates Piezo1 and interleukin-6 signaling to promote fibroblast activation"

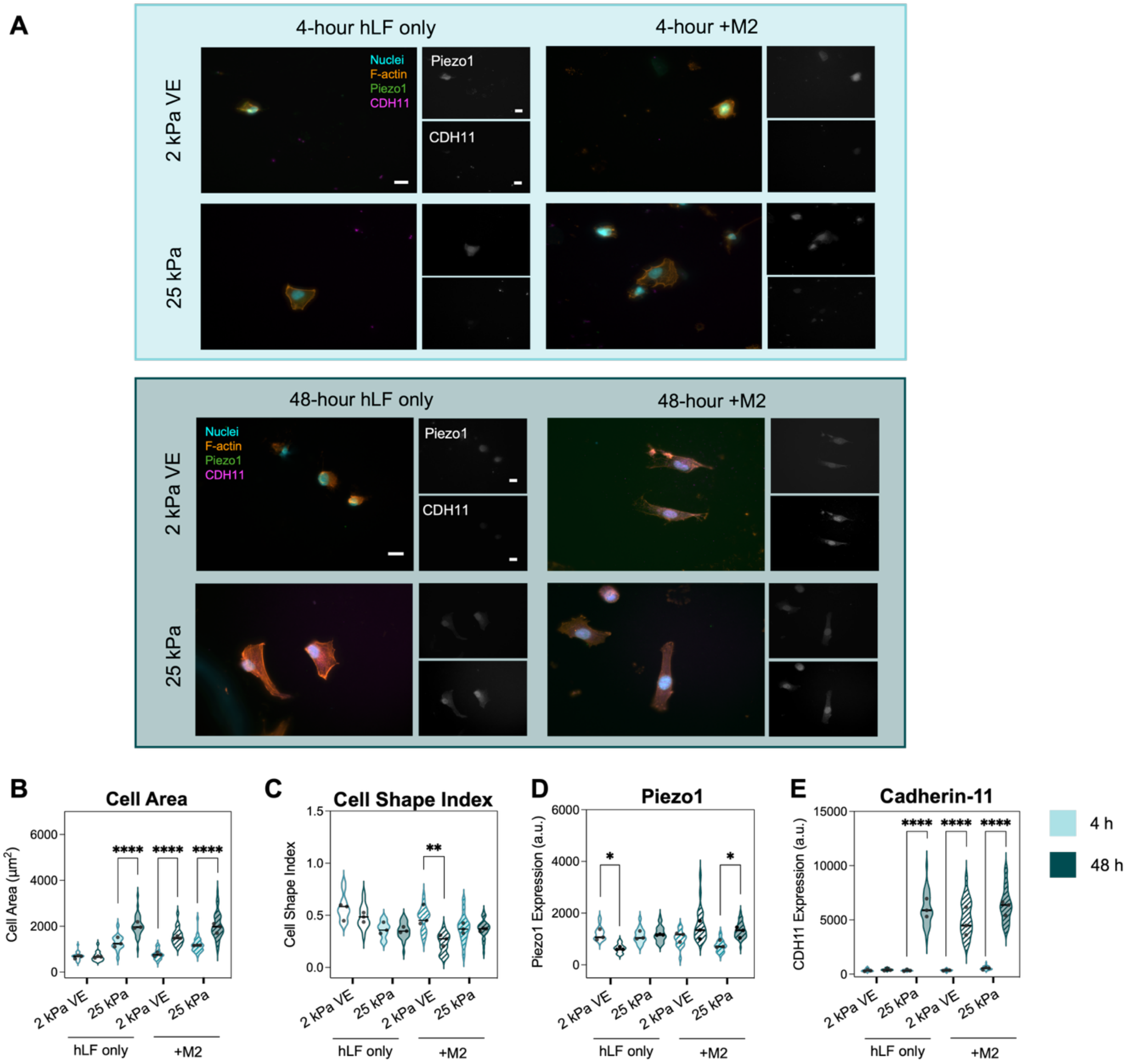


**Figure S1: Fibroblast spreading and CDH11 expression increase over 48 hours of culture, while Piezo1 levels remain consistent.** A) Representative images of human lung fibroblasts (hLFs) seeded on 2 kPa viscoelastic (VE) and 25 kPa elastic substrates with or without the addition of M2 macrophages after 4 hours (top panel) and 48 hours (bottom panel) of culture. Scale bars = 20 μm. B) WT fibroblasts cultured for 4 hours exhibited increased cell areas on 25 kPa hydrogels, while fibroblasts cultured for 48 hours also showed increased spreading on 2 kPa substrates in M2 macrophage co-cultures. Furthermore, fibroblasts cultured for 48 hours exhibited significantly higher areas compared to those cultured for 4 hours in all groups other than fibroblast only cultures on 2 kPa VE hydrogels, indicating that fibroblasts become more activated over time. C) Fibroblast morphology was similar between 4 and 48 hours of culture. D) Fibroblasts expressed consistent Piezo1 levels between 4 and 48 hours of culture. E) Fibroblasts cultured for 48 hours had significantly higher levels of CDH11 expression on 25 kPa hydrogels and in macrophage co-culture, independent of hydrogel stiffness, suggesting CDH11 may play a role in the increased activation observed between 4 and 48-hour cultures. Each point represents one hydrogel average, $n = 3$ hydrogels per group, 45-104 individual cells per group. Statistical analyses performed via two-way ANOVA with Tukey's HSD post hoc testing. $****p < 0.0001$, $**p < 0.01$, $*p < 0.05$.

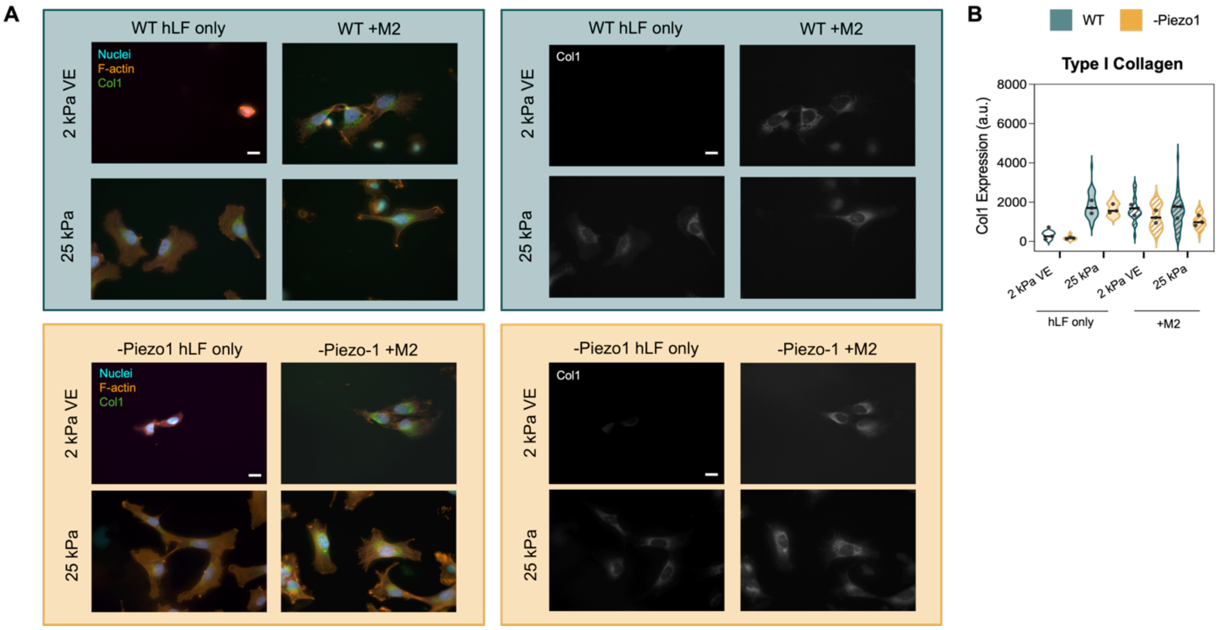


**Figure S2: Piezo1 inhibition does not significantly change type I collagen expression.** A) Representative images of wild type (WT) fibroblasts (top panel) and Piezo1 inhibited (-Piezo1) fibroblasts (bottom panel) seeded on 2 kPa viscoelastic (VE) and 25 kPa elastic substrates after 48 hours of culture. Scale bars = 20 μm. B) WT and Piezo1 inhibited fibroblasts exhibited increased expression of type I collagen on 25 kPa hydrogels and in co-culture with M2 macrophages regardless of hydrogel stiffness, with no significant differences between control and treatment groups. Each point represents one hydrogel average, $n$ = 3 hydrogels per group, 51-95 individual cells per group. Statistical analyses performed via two-way ANOVA with Tukey's HSD post hoc testing.

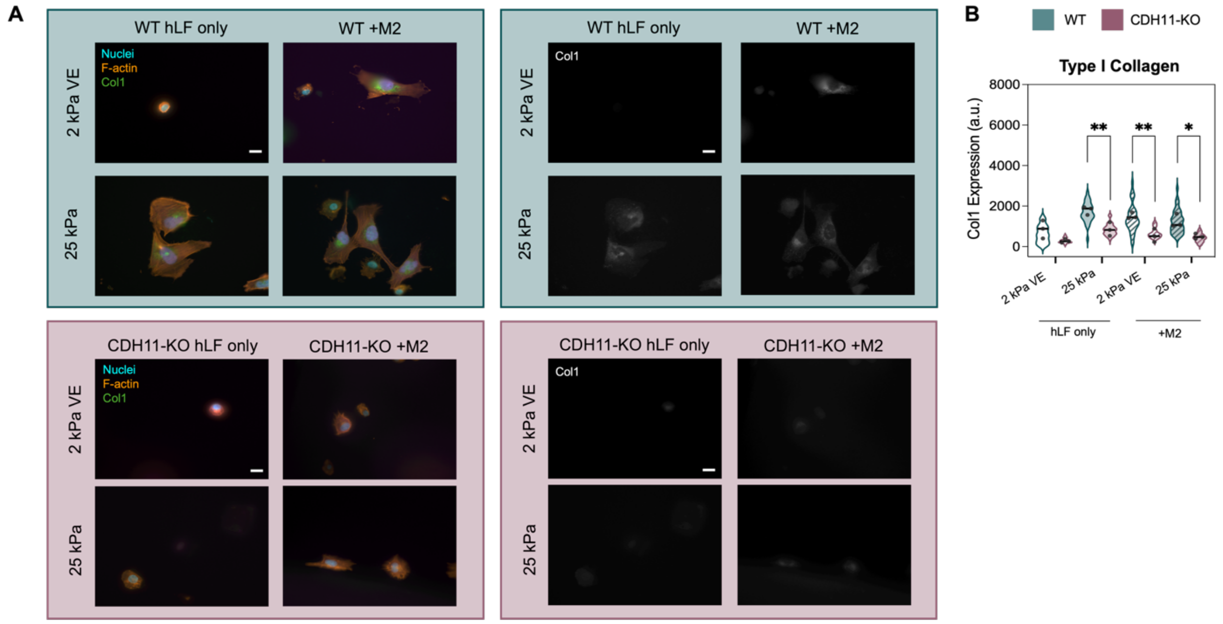


**Figure S3: Cadherin-11 knockdown reduces type I collagen expression in fibroblasts.** A) Representative images of wild type (WT) fibroblasts (top panel) and cadherin-11-knockout (CDH11-KO) fibroblasts (bottom panel) seeded on 2 kPa viscoelastic (VE) and 25 kPa elastic substrates after 48 hours of culture. Scale bars = 20 μm. B) WT fibroblasts exhibited increased expression of type I collagen on 25 kPa hydrogels and in co-culture with M2 macrophages regardless of hydrogel stiffness. However, CDH11-KO fibroblasts showed significantly reduced expression of type I collagen in these conditions, indicating a lack of response to mechanical or macrophage-derived cues. Each point represents one hydrogel average, $n = 3$ hydrogels per group, 52-115 individual cells per group. Statistical analyses performed via two-way ANOVA with Tukey's HSD post hoc testing. $**p < 0.01$, $*p < 0.05$.

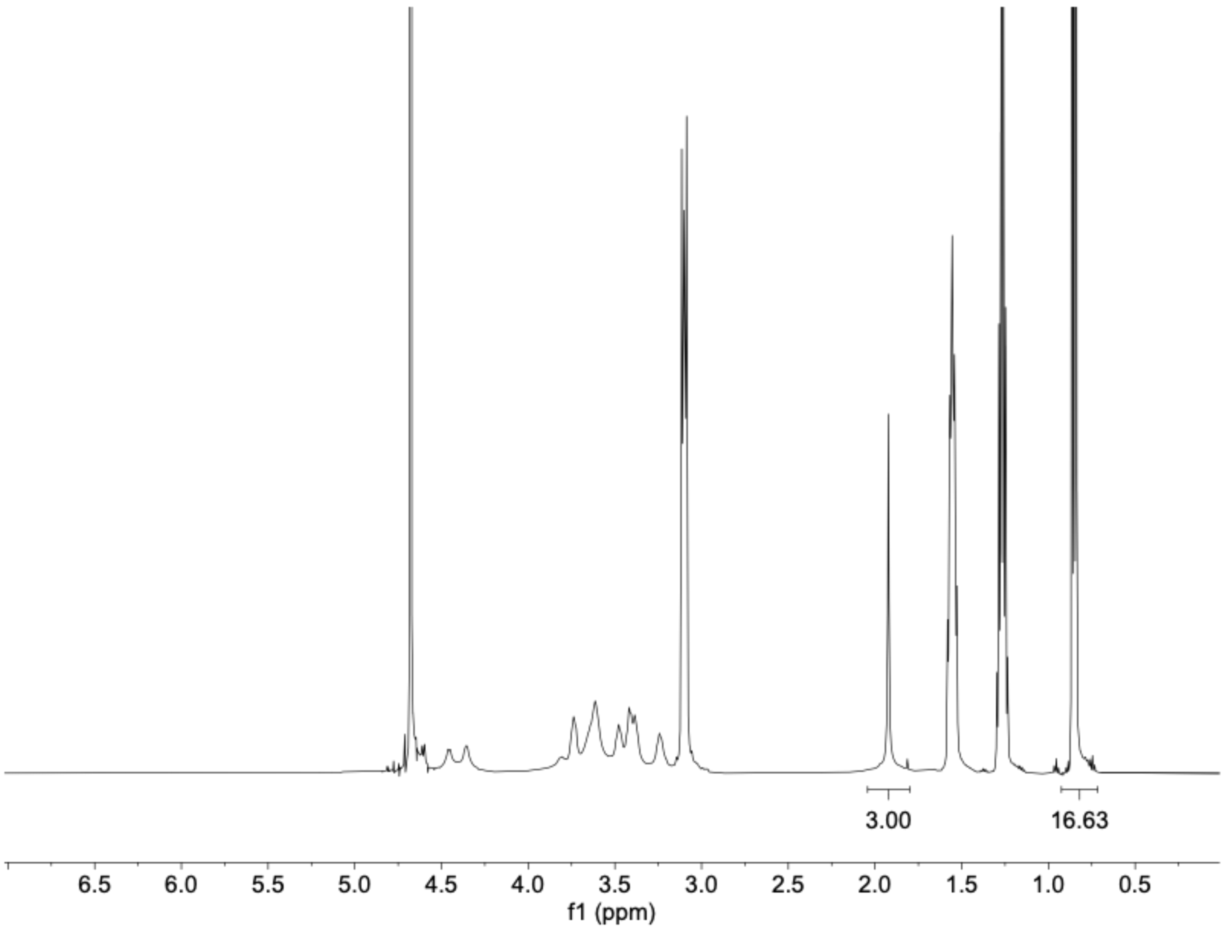


**Figure S4: $^{1}$H NMR spectra of tetrabutyl ammonium salt of hyaluronic acid (HA-TBA).** HA modification with TBA salt is determined by the integration of the TBA methyl-groups relative to the *N*-acetyl group of HA.

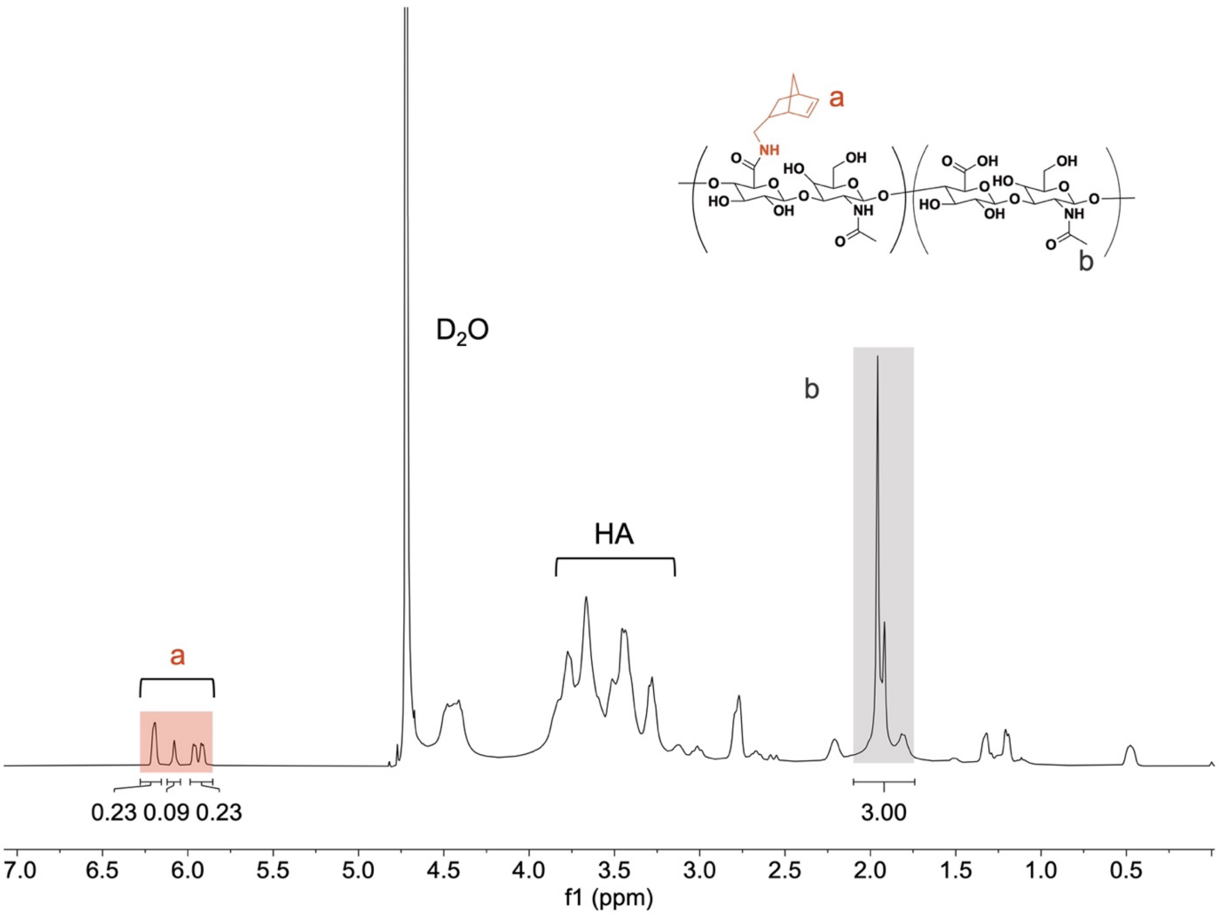


**Figure S5: $^{1}$H NMR spectra of norbornene-modified hyaluronic acid (NorHA).** Modification of HA with pendant norbornenes was determined to be 28%, indicated by the integration of peaks at $\delta$ = 5.95, 6.05, and 6.2 ppm (2H, ‘a’) normalized to the *N*-acetyl on HA (3H, ‘b’).

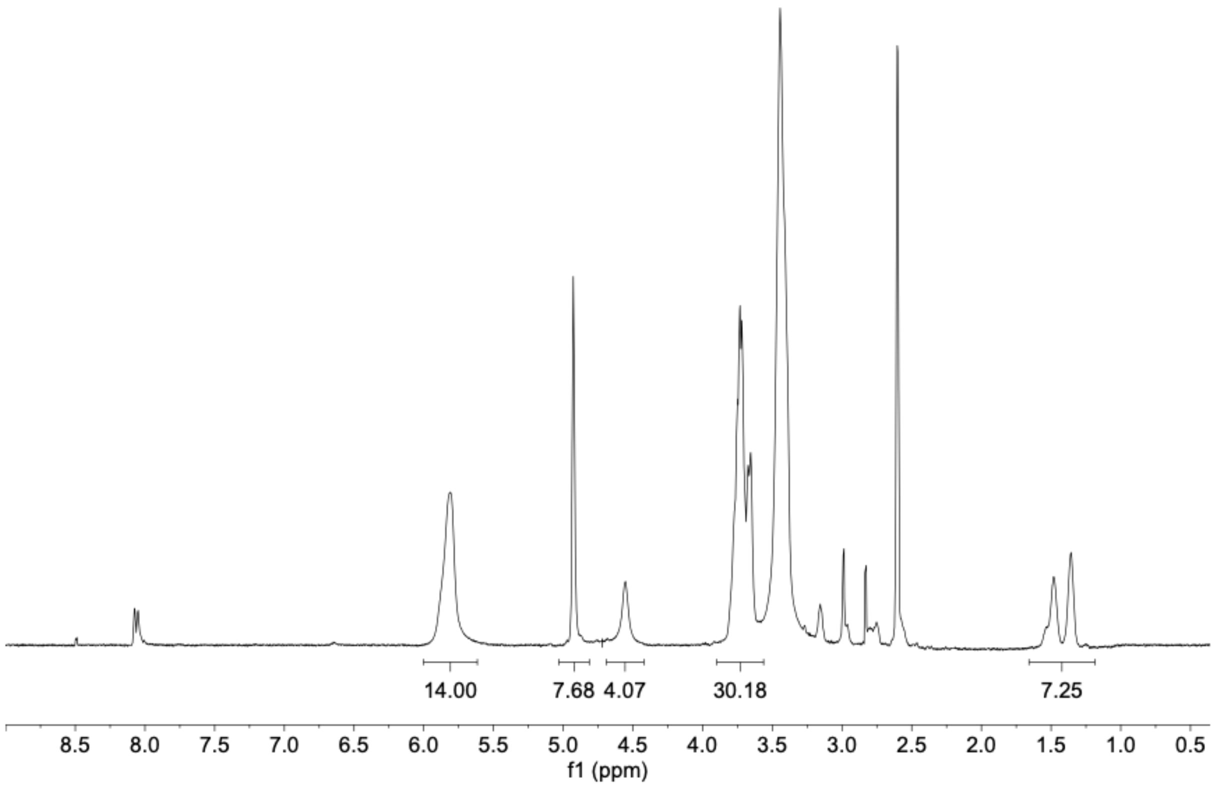


**Figure S6: $^{1}$H NMR spectra of 6-(6-aminohexyl)amino-6-deoxy-β-cyclodextrin (CD-HDA).** Modification of β-CD with HDA was determined to be 60%, indicated by the integration of peaks at $\delta$ = 1.14-1.6 ppm (12H).

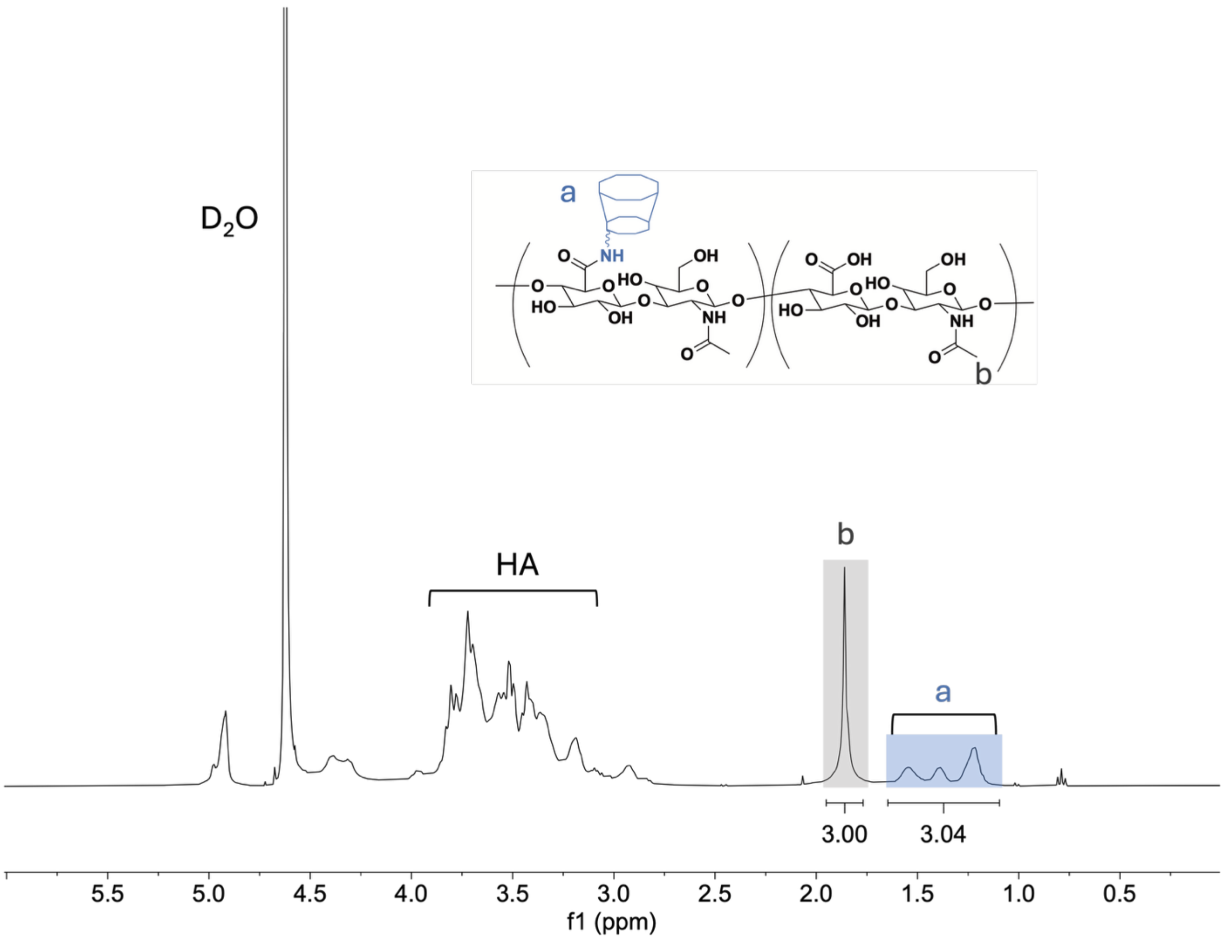


**Figure S7: $^1$H NMR spectra of β-cyclodextrin modified hyaluronic acid (β-CD-HA).** Modification of HA with pendant cyclodextrins was determined to be 25% by integration of hexane linker peaks at $\delta$ = 1.2-1.7 ppm (12H, a) relative to the *N*-acetyl group of HA (3H, b).

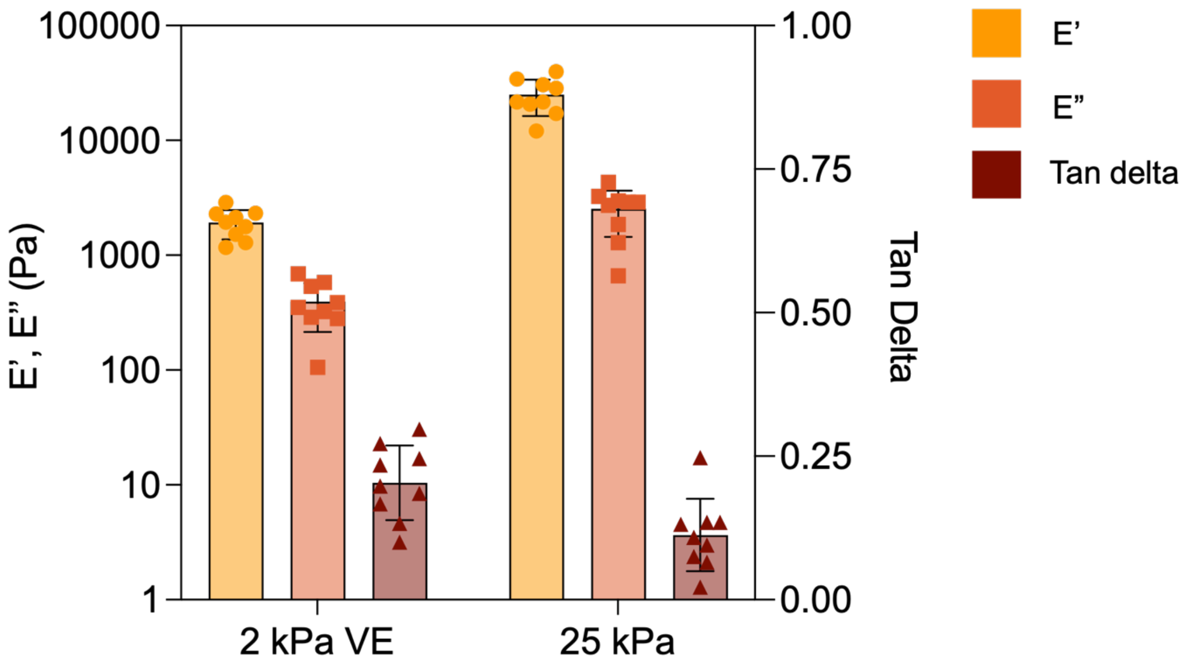


**Figure S8: Mechanical characterization of 2 kPa viscoelastic (VE) and 25 kPa hydrogels via nanoindentation.** Storage (E′) moduli, loss (E″) moduli, and tan delta (E"/E') were determined using dynamic mechanical analysis (DMA) at a frequency of 1 Hz at a depth of 4 µm at 3 points per hydrogel replicate. *n* = 3 hydrogels per group.
