## Supplemental File 1 for "Cadherin-11 integrates Piezo1 and interleukin-6 signaling to promote fibroblast activation"

#### Untreated

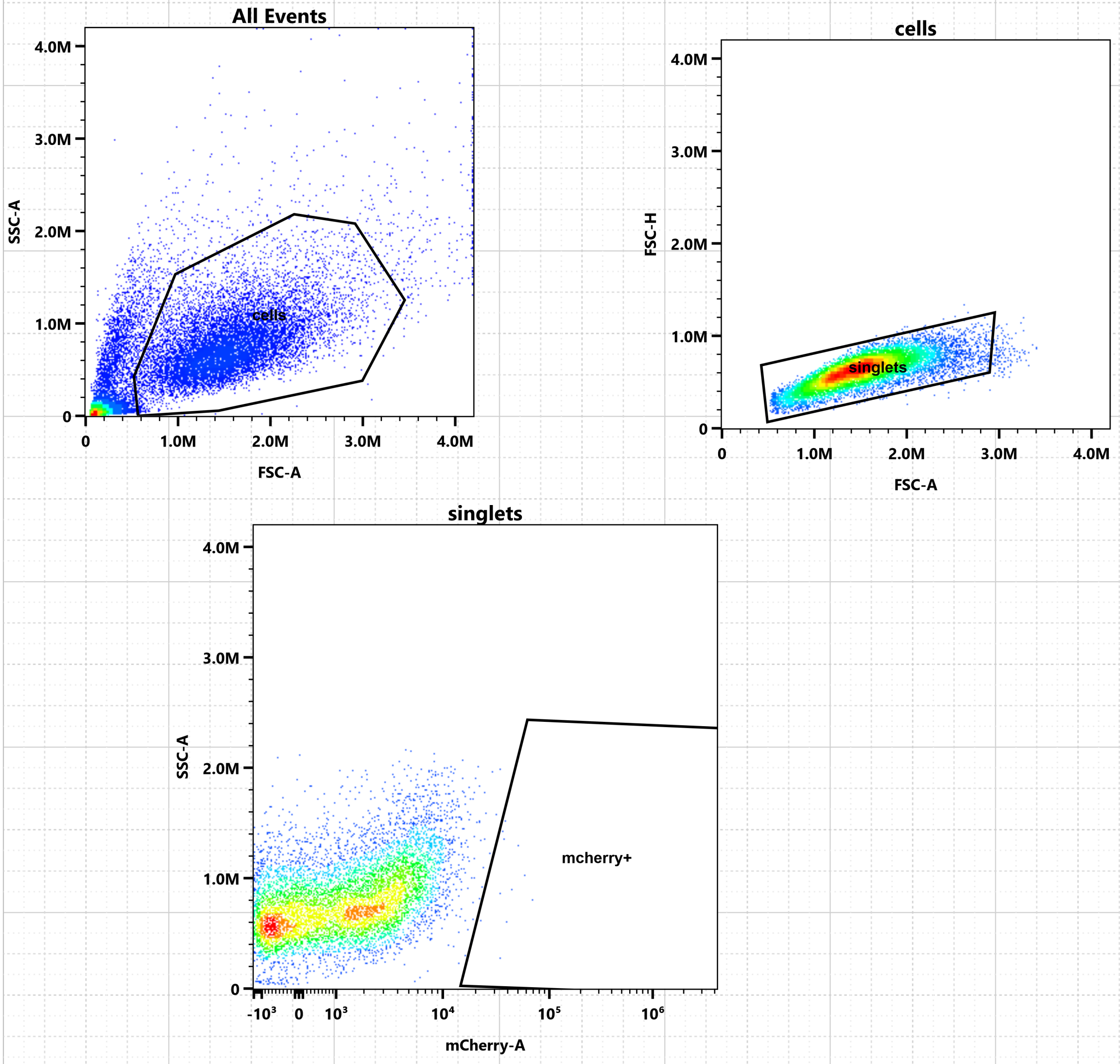

All Events
SSC-A
FSC-A
4.0M
3.0M
2.0M
1.0M
0
1.0M
2.0M
3.0M
4.0M
cells
FSC-H
FSC-A
singlets
SSC-A
mCherry-A
mcherry+
-10³
0
10³
10⁴
10⁵
10⁶

#### Untreated

##### Population Hierarchy
##### 3.22.24 Fibroblasts-Group_001-Unstained

| Population | % Total | Count |
|---|---|---|
| All Events | 100.00 | 25,000 |
| cells | 45.69 | 11,423 |
| singlets | 45.00 | 11,249 |
| mcherry+ | 0.04 | 11 |

### Transfected

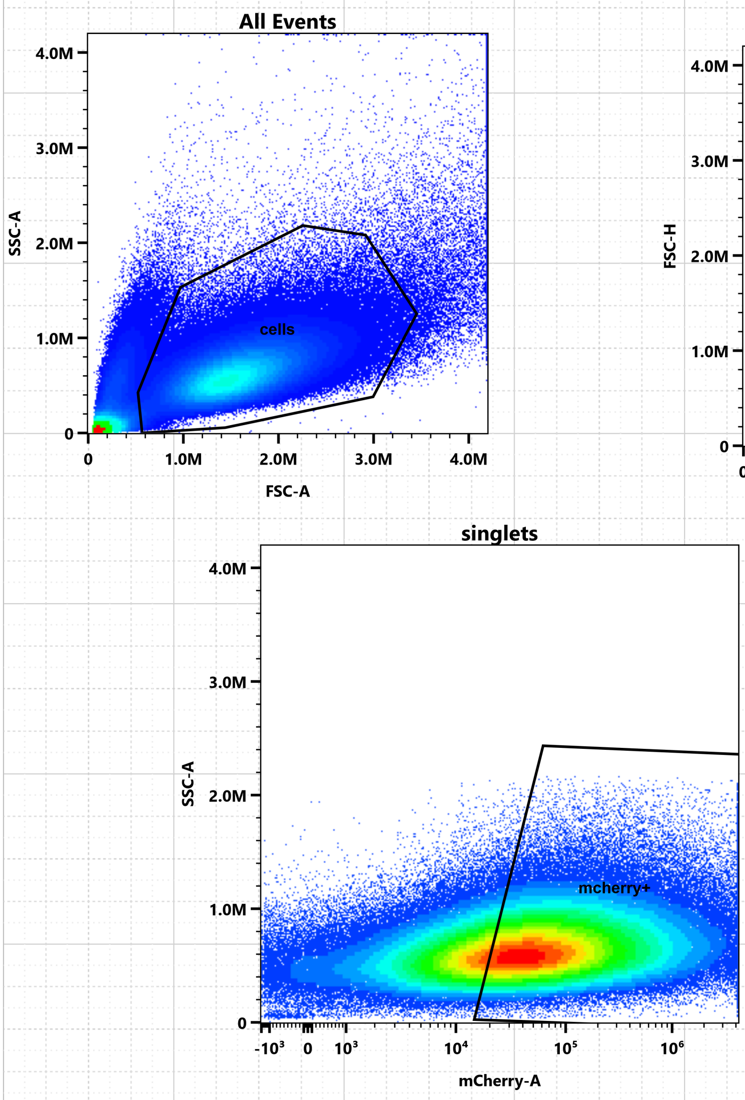

#### Transfected

Population Hierarchy
3.22.24 Fibroblasts-Group_001-3.75

| Population | % Total | Count |
|---|---|---|
| All Events | 100.00 | 500,000 |
| cells | 64.87 | 324,363 |
| singlets | 62.89 | 314,436 |
| mcherry+ | 39.46 | 197,304 |
